# The Horizontal Ladder Test (HLT) protocol: A novel, optimized, and reliable means of assessing motor coordination in *Sus scrofa domesticus*

**DOI:** 10.1101/2023.12.13.571517

**Authors:** Xiaobo Liu, Ana G. Gutierrez, Arlette Vega, Joshua O. Willms, Jackson Driskill, Praneetha Panthagani, Jordan Sanchez, Monica Aguilera, Brittany Backus, Jeremy D. Bailoo, Susan E. Bergeson

## Abstract

Pigs can be an important model for preclinical biological research, including neurological diseases such as Alcohol Use Disorder. Such research often involves longitudinal assessment of changes in motor coordination as the disease or disorder progresses. Current motor coordination tests in pigs are derived from behavioral assessments in rodents and lack critical aspects of face and construct validity. While such tests may permit for the comparison of experimental results to rodents, a lack of validation studies of such tests in the pig itself may preclude the drawing of meaningful conclusions. Here, we present a novel, optimized, and reliable horizontal ladder test (HLT) test protocol for evaluating motor coordination in pigs and an initial validation of its construct validity using voluntary alcohol consumption as an experimental manipulation.

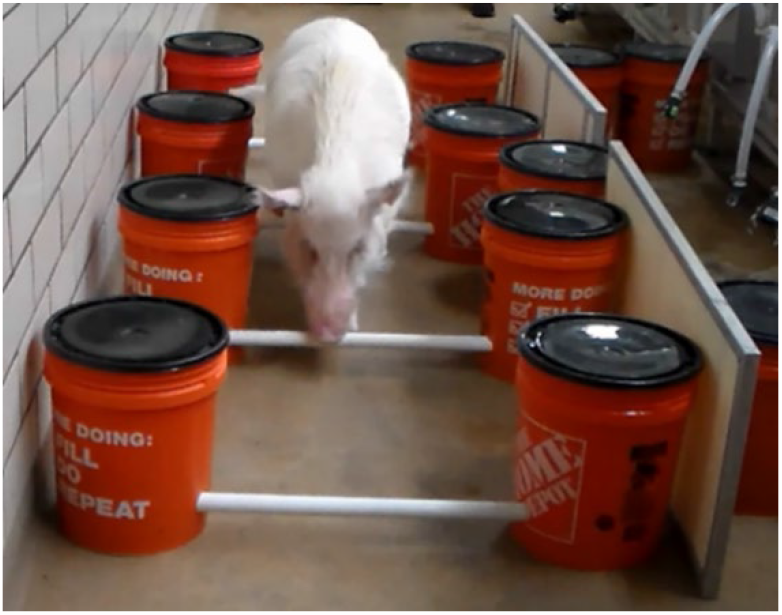

## INTRODUCTION

While rodents are the most prevalently used animal in preclinical animal research, high rates of translational failure concerning drug development have brought into sharp focus the need to study mammalian species that are physiologically more similar to humans, particularly in relation to the aspects of the diseases being modeled^1-8^. Pigs are an important model in preclinical biomedical research, historically accounting for approximately 6% of all USDA species protected under the Animal Welfare Act^8^. Pigs share numerous physiological and neuroanatomical similarities with humans^9,10^ and consequentially are used in the research setting for the study of oncology, cardiovascular diseases, pediatric disorders, regenerative medicine, transplantation, medical imaging, genomic and reproductive biotechnology, neurological diseases, surgical innovation, gene therapy/immunotherapy, infectious diseases, effects of microbiota, metabolism, nutrition, and education^11,12^.

There is a relative dearth of reliable, validated means of behavioral assessments for pigs within a biomedical setting^13,14^. One of the commonly evaluated behavioral phenotypes in the laboratory pig is impaired motor coordination. The apparatuses used to evaluate motor coordination differences in pigs includes the open field, the balance/inclined beam, and the treadmill. It has been argued that analysis of pig behavior within these apparatuses permits researchers to evaluate motor behavior within a controlled environment. However, these behavioral assessments are derived from rodent models with little emphasis on the degree to which rodents and pigs differ and the consequences of such differences on the validity of derived experimental results. For example, the inclined beam test, which is very similar to the balance beam test for rodents, can also be potentially dangerous to a pig with impaired motor or cognitive function if it were to fall.

Moreover, at baseline/unimpaired levels, some animals cannot walk at least halfway up the beam, calling into question the reliability and validity of such assessments^15^. Similarly, in the open-field test, the ability to measure locomotor behavior while inducing conflict in the emotional/arousal domains of behavior often yields conflicting results across replicate studies^13,14^. Additionally, the typical open field arena for pigs is orders of magnitude smaller than the typical foraging distance observed within an ethological setting, calling into question the ethological validity of this test^17^. Assessment of gait using a treadmill often requires advanced equipment and facilities, which are generally cost-prohibitive and for which infrastructure is sorely lacking^16,17^. Therefore, inexpensive apparatuses that are safe, easy-to-train, and which produce reliable and repeatable evaluations of motor behavior are critically needed for assessing motor impairment in pigs.

To address this knowledge gap, an apparatus modeled after a horizontally placed ladder that the animal had to cross was designed. Given that pigs have short legs, considerable weight, and an ungainly gait^11,12,14,18^, this approach involved tailoring the protocol to minimize physical contact with the apparatus–the operational definition of a motor coordination error used here. The apparatus design was modular, such that the rung height could be readily adjusted from 3 to 6 inches, in 1-inch increments. It was hypothesized that: 1) By increasing the rung height to 6 inches during the initial training phase, the pigs would be forced to lift their feet in order to be able to cross the ladder, and 2) The pigs would make significantly fewer errors as the rung height was lowered from 6 to 3 inches, in 1-inch increments across training sessions.

As an initial validation of construct validity, changes in motor coordination as a consequence of voluntary consumption of escalating concentrations of alcohol (ethanol) in water was evaluated. It was predicted that, similarly to humans, as the percentage of ethanol in water consumed increased from 2.5 to 10% during voluntary drinking sessions, a significantly greater number of motor coordination errors would be observed when performing the HLT task.

## PROTOCOL

### 1. Animal husbandry

Five female Sinclair miniature pigs purchased from Sinclair BioResources (Auxvasse, MO) at seven months of age were used in this study. Before entering the experiment, the pigs were acclimated to the local laboratory environment and individually housed for two months. Animals were housed in an environmentally controlled vivarium with an 11/13 light/dark cycle, with lights off at 18:00. The temperature in the housing room was maintained at 23°C – 25°C, with humidity levels set to 30% – 60%. The Laboratory Animal Resources Center staff provided the pigs with two meals daily, at 08:00 and 13:30, during weekdays, while a single meal was given at 08:00 on weekends, with twice the usual weekday food quantity allotment. Access to filtered tap water was unrestricted for the pigs, except during specific two-bucket, free-choice alcohol access testing periods when house water was temporarily turned off (see Section 3 below). All experimental procedures conducted in this study were approved by the Institutional Animal Care and Use Committee at Texas Tech Health Sciences Center.

### 2. The Horizontal Ladder Test-Description and Protocol

#### 2.1 Apparatus Description

The horizontal ladder is comprised of four rungs and eight 5-gallon buckets. The rungs of the horizontal ladder apparatus are comprised of lengths of Schedule 40 polyvinyl chloride (PVC) tubes, 1.5 inches in diameter and 88.5 inches in length (to accommodate the width of the largest pig used in our study). The length of PVC tubes used can be increased or shortened depending on the size of the animal in a given study (Figure 1A). Each rung is capable of being slotted into 5-gallon buckets at a height of 3, 4, 5, or 6 inches (Figure 1B) from the floor through 1.5-inch diameter drilled holes. At most, the fall distance for a pig is 6 inches, minimizing, if not essentially eliminating, the potential for injury. Each bucket is weighted with 35 lbs. sandbags for stability and to minimize the deviation of our pre-specified rung distance interval of 29.5 inches (the maximum length of our pigs). This distance can be adjusted as needed (see 2.2 below). The total cost of the apparatus to build, using materials readily available at a big-box hardware store, was less than $250.00 excluding the sand, which can be substituted for whatever weight sources that are handy (see Table 1 for details).

**Table 1.**
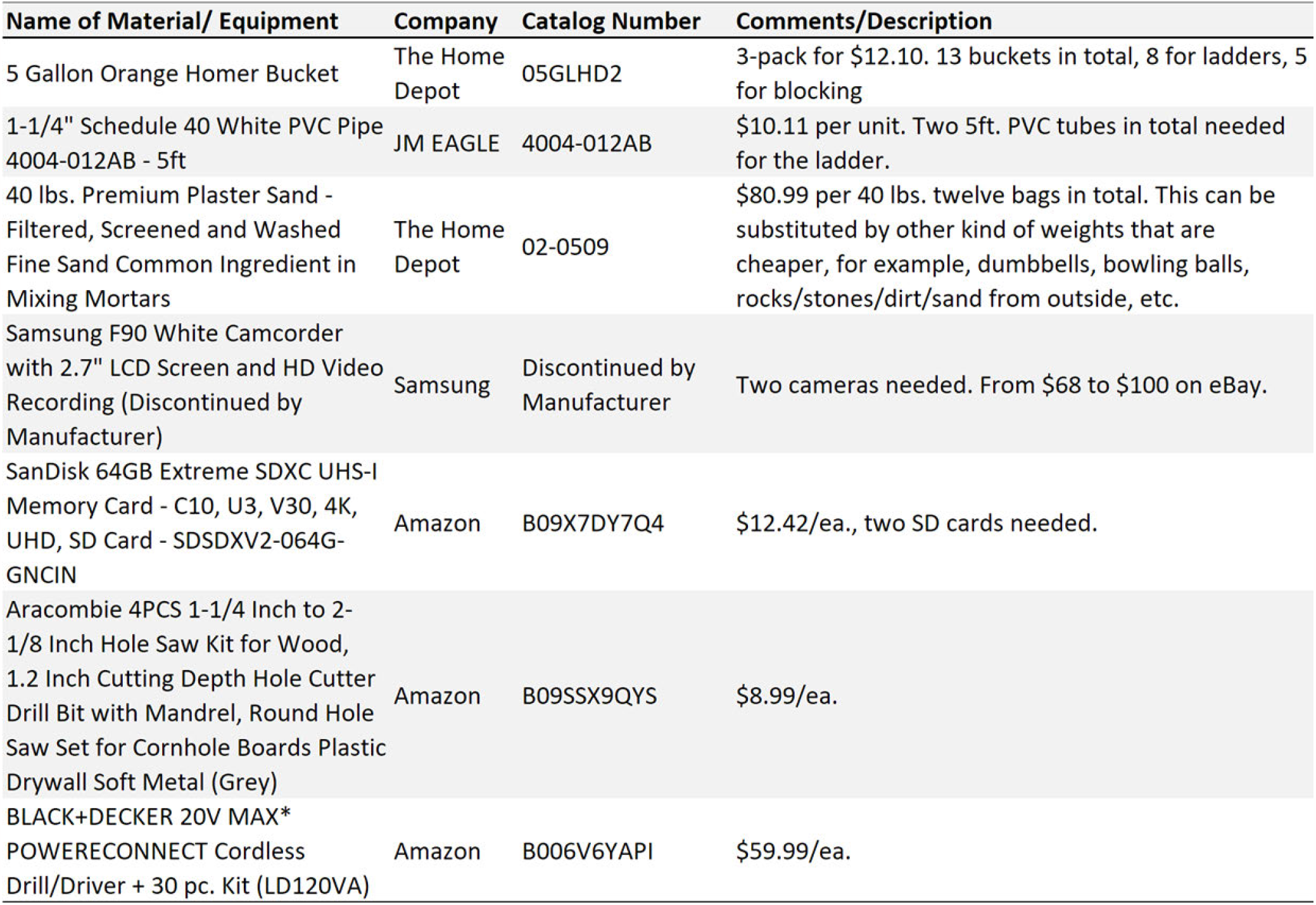
List of materials used to construct the HLT

**Figure 1.**
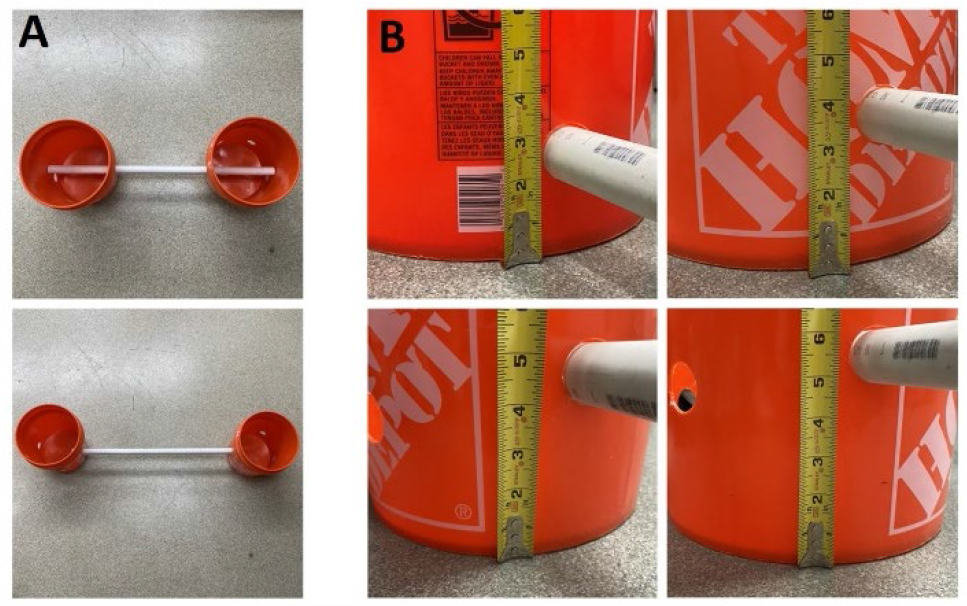
Example of the adjustable length and height of the rung. (A) The upper picture shows the shortest length of the rung, and the lower picture shows the longest length of the rung. (B) From the top left to the bottom right, the rung heights were at 3 in, 4 in, 5 in, and 6 inches with holes drilled with spaces around each bucket.

#### 2.2 Apparatus Setup

1. Place the rungs of the ladder into each 5-gallon weighted bucket at either 3, 4, 5, or 6 inches, depending on the training day. Attach the other end of the PVC tube to another bucket so that each end of the rung is completely within the bucket.
2. Arrange the assembled buckets in a parallel fashion on the floor, with a gap of 29.5 inches—or the width of your largest pig—between them. Laboratory tape may be used on the floor to designate this distance and for easy replacement of the buckets should they be moved. The width of the tubes can be easily adjusted by sliding the tube more or less into the bucket (± 10 inches in each bucket, 20 inches total) to accommodate the average width of each pig. The sides of the ladder can be blocked using easily obtainable materials, such as additional weighted buckets, or by placing one side of the ladder against a wall. The accommodations effectively ensure that the animal is only able to traverse the ladder by proceeding forward or backward across the ladder, with no opportunity for exiting the ladder until the last rung is reached.
3. Set up the tripods and video cameras, one on each end of the ladder, such that the view of the animal is unobstructed. Ideally, the cameras should record video in a minimum resolution of 720p and with stereo audio. Refer to Figure 2 for representative images of the assembled ladder and setup of the camera viewpoints.
4. Position one investigator at the far end of the ladder while a second investigator stands at the opposite end of the ladder. Each investigator can encourage the animal to cross the ladder using food rewards (e.g., marshmallows, breakfast cereals, etc.)

**Figure 2.**
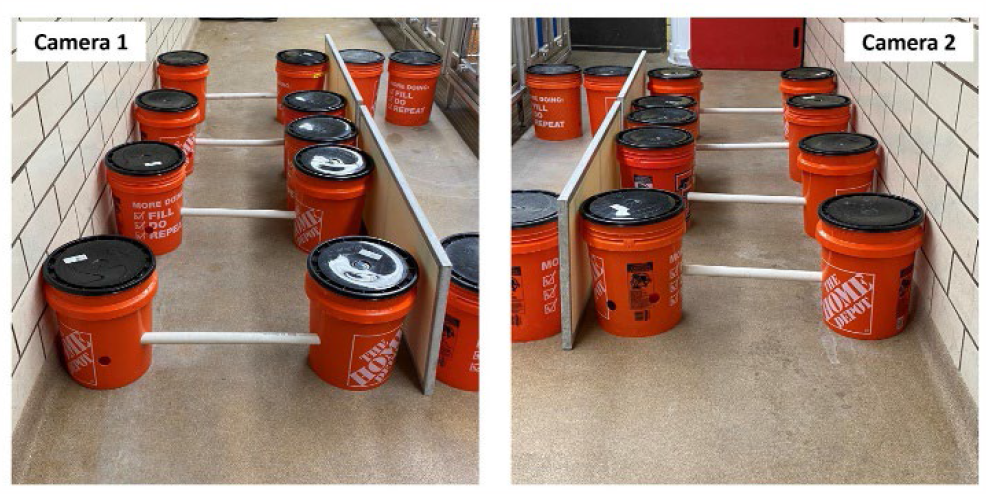
The horizontal ladder setup with exemplars of the camera-view achieved from both sides.

#### 2.3 Behavioral Training

Training to cross the ladder proceeds in a stepwise process, which minimizes the number of things that the animal has to learn in a single step, and which also minimizes the potential for off-task behavior. Such a stepwise process also permits re-training at a specific step if recalcitrant behavior is observed.

*<u>Step 1</u>*. An investigator guides each pig to cross the ladder, offering treats as a reward whenever a pig completes a single pass of the ladder. During this step, the rungs are set to the highest level of 6 inches, so that the animal is forced to lift its legs in order to cross.

*<u>Step 2</u>*. Here, animals are only rewarded when they complete two passes of the ladder, with the rung height set to 6 inches, before returning to its initial starting location. The second investigator assists in turning the animal around once the animal has completed its first pass. This step continues until the animal can, at a minimum, make 4 continuous double crossings of the ladder. If significant issues are observed (uncommon), re-train the animal on Step 1.

*<u>Step 3</u>*. Here, animals are only rewarded when they complete two passes of the ladder, with the rung height set to 5 inches, before returning to their initial starting location. The second investigator assists in turning the animal around once the animal has completed its first pass. This step continues until the animal can, at a minimum, make 4 continuous double crossings of the ladder. If off-task behavior is observed (e.g., refusal to cross), re-train the animal on Step 2.

*<u>Step 4</u>*. Here, animals are only rewarded when they complete two passes of the ladder, with the rung height set to 4 inches, before returning to their initial starting location. The second investigator assists in turning the animal around once the animal has completed its first pass. This step continues until the animal can, at a minimum, make 4 continuous double crossings of the ladder. If off-task behavior is observed (e.g., refusal to cross), re-train the animal on Step 3.

*<u>Step 5</u>*. Here, animals are only rewarded when they complete two passes of the ladder, with the rung height set to 3 inches, before returning to their initial starting location. The second investigator assists in turning the animal around once the animal has completed its first pass. This step continues until the animal can, at a minimum, make 4 continuous double crossings of the ladder. If off-task behavior is observed (e.g., refusal to cross), re-train the animal on Step 4. Training is considered complete, when the pigs consistently score less than 4 points during a “single” crossing at a rung height of 3-inches (see Table 2).

**Table 2.**
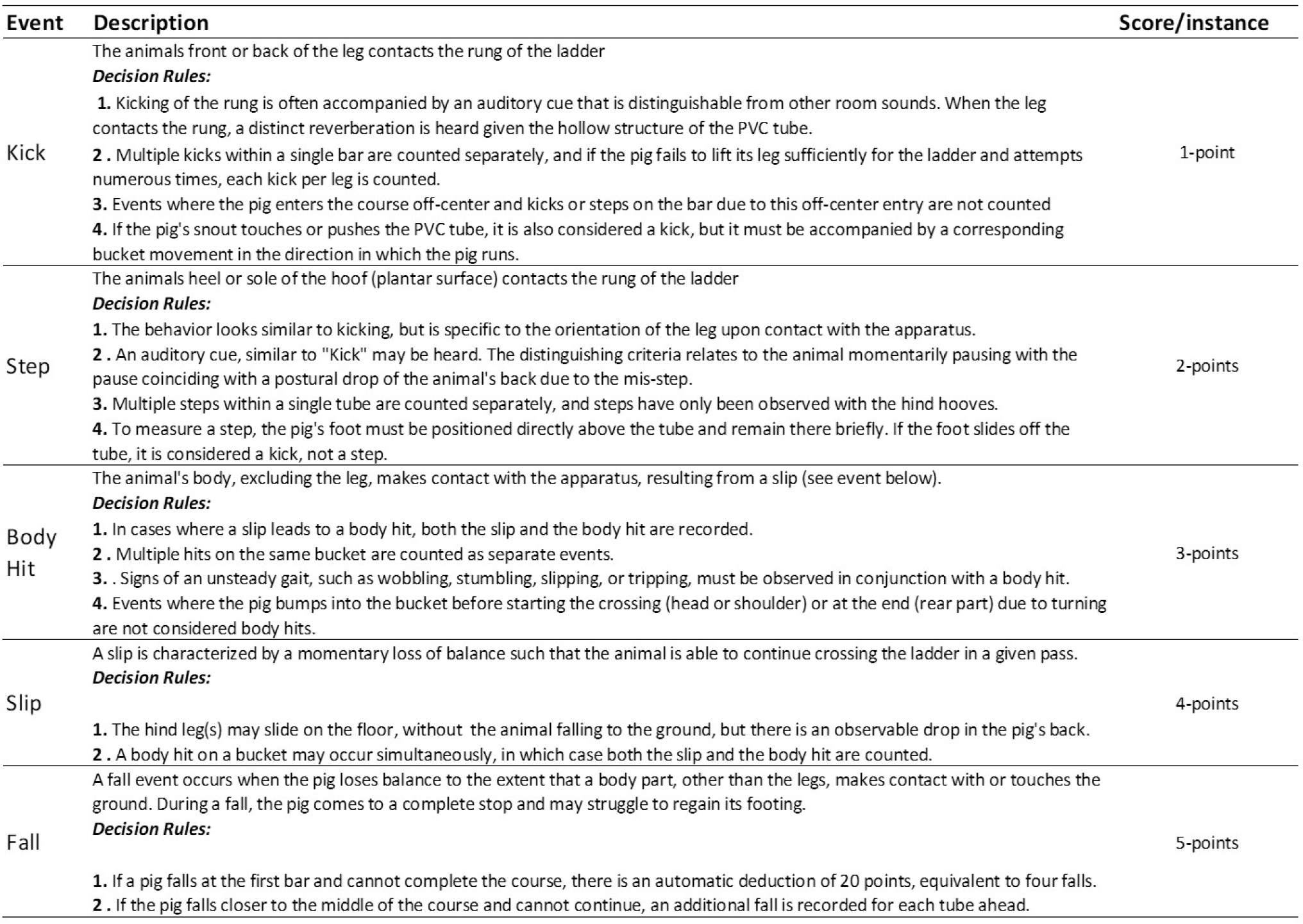
Ethogram for the scoring of motor coordination in the HLT.

#### 2.4 Video Scoring

##### 2.4.1 Video data preparation

Videos should be screened for excessive noise, completeness (e.g., equipment failure), and other technical issues (e.g., improper placement of cameras). Only complete data for each pig should be analyzed. In cases where animals belong to experimental groups (e.g., treated vs control), videos can be renamed so that the experimenter is blinded to treatment. It may not always be possible to blind the experimenter to treatment if, for example, the hair/bristle of the pigs are different colors. Where possible, the experimenters who test the animals should be different from those who code the video data to avoid bias.

##### 2.4.2 Behavioral Ethogram

An ethogram was developed for the scoring of errors in motor coordination that can be generalized to other laboratory/vivarium settings. The ethogram focused on five observable and objective criteria: kicking (of the rungs or buckets), stepping (on the rungs), body contact with the apparatus (rungs or buckets), slipping (momentary loss of balance), and fall (complete loss of balance with contact of the body unto the floor) (c.f., Table 2 and Supplementary Videos). Data were scored using the open-source software Behavioral Observation Research Interactive Software (BORIS), using the previously established methods for reliability assessment^19-24^. For these experiments, inter- and intra-rater reliability was established at 80% concordance. The primary outcome measure from these evaluations was the total score, at each rung height, across all passes. Here, higher scores reflect a greater number of errors and vice versa.

##### 2.4.3 Statistical Analyses

Depending on the sample size and adherence to distributional assumptions, a within-subject ANOVA may be used for longitudinal analysis of changes in motor coordination behavior. For these data, a non-parametric Friedman test for analysis of repeated measures data was used, followed by *post hoc* corrected Dunn’s test.

### 3. Evaluation of construct validity using voluntary ethanol consumption

#### 3.1 The two-choice bucket test

The pigs were given a choice between two buckets: one containing water and the other with increasing alcohol (ethanol) concentrations, starting at 2.5% and progressing to 5%, then 7.5%, over two weeks for each concentration. Subsequently, the ethanol concentration was elevated to 10%, remaining at this level for eight weeks. The location of water and ethanol buckets (left or right side) was alternated daily to control for side-biased responding. Animals had access to alcohol from 10:00 to 17:00 each day, with a maximum volume limit of 3.5 liters, aiming to prevent discontinuous alcohol consumption or any potential life-threatening situations arising from the adverse effects of excessive alcohol intake. After 17:00, access to house water was reinstated until the next round of alcohol exposure.

#### 3.2 Evaluation of motor coordination errors

Before collecting baseline data, animals were trained to cross the ladder using the methods described in Section 2. Baseline data, at a rung height of 3 and 6 inches, was then collected one week prior to the commencement of alcohol exposure. The animals were then evaluated for motor coordination errors in the HLT at a rung height of 3 and 6 inches for each ethanol concentration, and the data was compared to baseline levels.

#### 3.3 Blood alcohol concentration (BAC) determination

Blood samples were obtained from the pigs’ ear veins approximately one hour after they had concluded their daily drinking sessions for each alcohol concentration exposure. The BACs were then analyzed using Gas Chromatography (GC). Here, 50 μl of blood was placed into a GC vial along with 100 μl of water, and the vial cap was securely sealed to prevent any vapor leakage. Prior to GC analysis, these vials were stored on ice. For quality analysis and quality control (QAQC) evaluation, a series of standard solutions (0, 0.025, 0.5, 1, and 2 mg/mL) were analyzed with actual samples. The final BAC values were computed based on the resulting standard curve.

#### 3.4 Statistical Analyses

All statistical analyses were conducted in GraphPad Prism v10.0.3. A non-parametric Friedman test, for analysis of repeated measures data was used, followed by post hoc corrected Dunn’s test to evaluate motor coordination deficits in the HLT. For analysis of BAC, the median of each group was compared with the standard value for intoxication (0.8 mg/ml) using a one-sample t-test.

## REPRESENTATIVE RESULTS

### As predicted, the motor coordination score in the HLT decreased as the rung height was decreased from 6 to 3 inches

A sequential training approach was implemented as detailed in section 2 of the HLT protocol, where the rung height of the ladder gradually decreased from 6 inches to 3 inches. Figure 3 illustrates representative results of the final day of training at each specific rung height. Initially, the motor coordination score in the HLT was higher and more variable at 6 and 5 inches. However, as training progressed, each pig’s performance became more consistent, with lower variability and a lower score, at the 3-inch rung height. On the last training day at the 3-inch height, the median score was low, at 5 points from the total of six passes.

**Figure 3.**
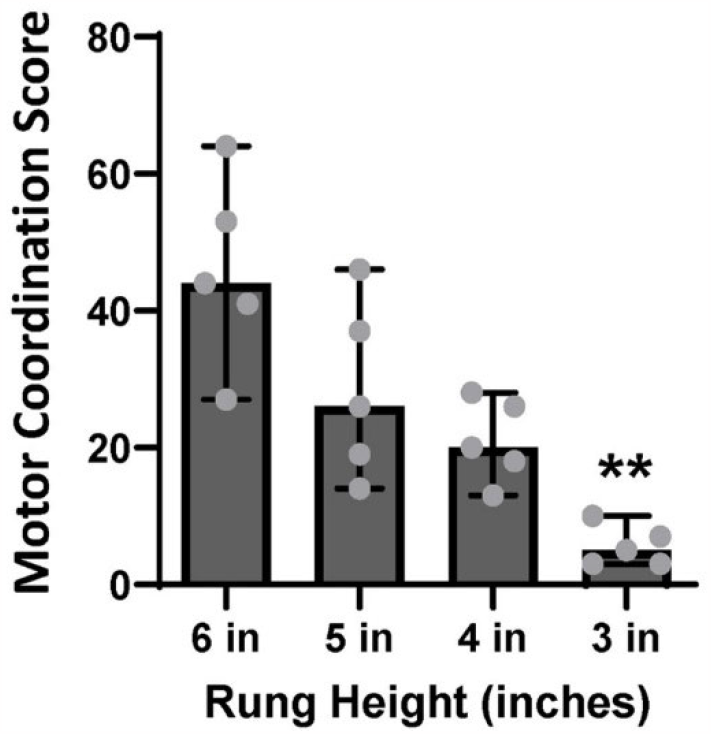
Motor coordination score on the last day of training at each height. As the rung height decreased across training, the motor coordination score significantly decreased (6 in vs. 3 in, p=0.0014) and was less variable (the animals performed significantly better at 3 inches in height). Bars represent medians ± 95% CI.

Since the training data were consistent with our predictions, the next procedure was to validate these results by evaluating motor coordination subsequent to voluntary alcohol consumption. This validation assessment occurred approximately one year after training was completed.

### The pigs drank a larger amount of alcohol as ethanol concentration increased and drank to intoxication levels

As shown in Figure 4A, a significant increase in ethanol consumption was observed at ethanol concentrations of 7.5% (p=0.0087) and 10% (p=0.0197), relative to 2.5% concentration levels. Notably, ethanol consumption (g/kg) was less variable at lower concentrations (2.5% and 5%) and more variable among individual pigs at higher ethanol concentrations (7.5% and 10%). This pattern of results is similarly reflected in the BAC data (Figure 4B).

**Figure 4.**
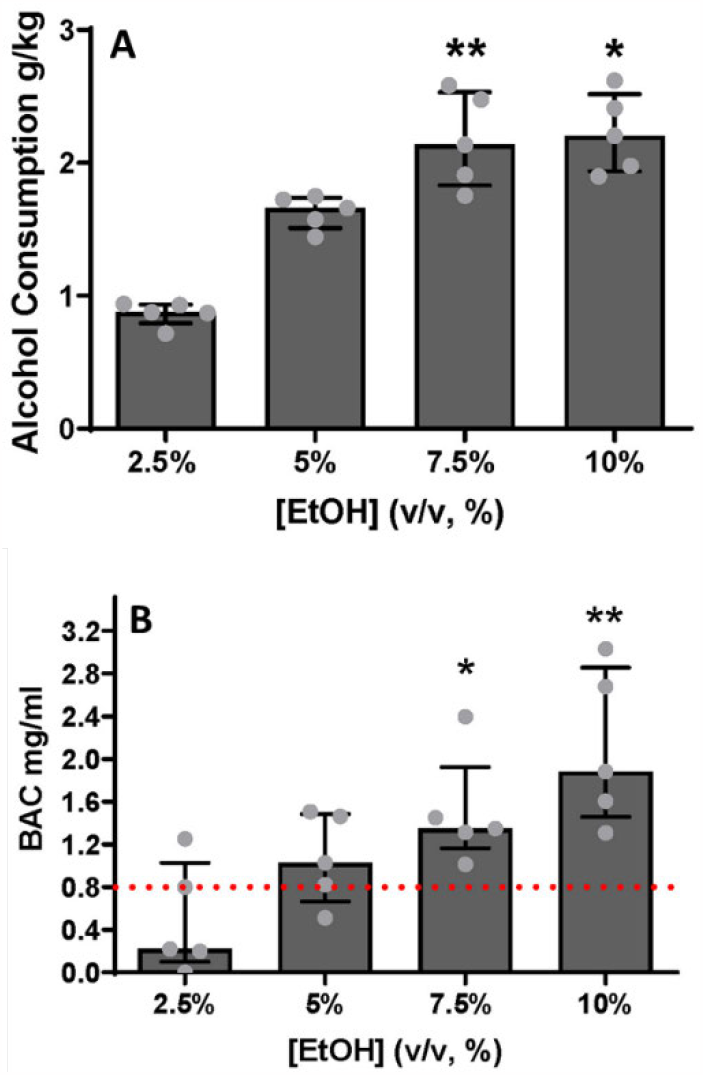
Alcohol consumption and blood alcohol concentration (BAC) increased across 2.5, 5, 7.5, and 10% alcohol concentrations. **A**. Animals consumed more alcohol, with increasing alcohol concentration, significantly escalating their intake at 7.5% (p=0.0087) and 10% (p=0.0197) relative to 2.5% levels. **B**. At 2.5% concentration levels, two pigs reached the intoxication level criterion defined by NIAAA, while at 5%, four pigs achieved criterion. All pigs reached intoxication criterion at 7.5% and 10%, with a statistically significant difference being observed compared to the standard value of intoxication (0.8 mg/ml) at 7.5% and 10% ethanol concentration (p= 0.0396 and p=0.0162, respectively). Bars represent medians ± 95% CI.

Two pigs drank to intoxicating levels, 0.8 mg/ml (0.08 g/dL), as defined by National Institute on Alcohol Abuse and Alcoholism (NIAAA), already at 2.5% ethanol concentration levels. As alcohol concentration increased, four pigs reached the intoxication criterion at 5% ethanol concentration levels, and all pigs reached criterion at 7.5 and 10% ethanol concentration levels (p=0.0396 and p=0.0162, respectively). The next step was to evaluate motor coordination deficits as a consequence of consuming alcohol to intoxicating levels.

### The number of motor coordination errors increased with increased ethanol consumption

All animals showed good recollection of the HLT task, one year subsequent to training, with the median score being lowest at baseline (no ethanol exposure, rung height fixed to 3 inches, Figure 5). As the concentration of ethanol increased, the score in the HLT also increased. Consistent with our BAC data, the highest scores were observed at 7.5% and 10% ethanol concentrations, with a statistically significant difference relative to baseline being observed at 10% ethanol concentration (p=0.0201). The same pattern of performance was also observed at 6 inches (the highest rung height), demonstrating that the difference in motor coordination score was not a consequence of the task being simpler to perform at a lower rung height of 3 inches (Figure 5).

**Figure 5.**
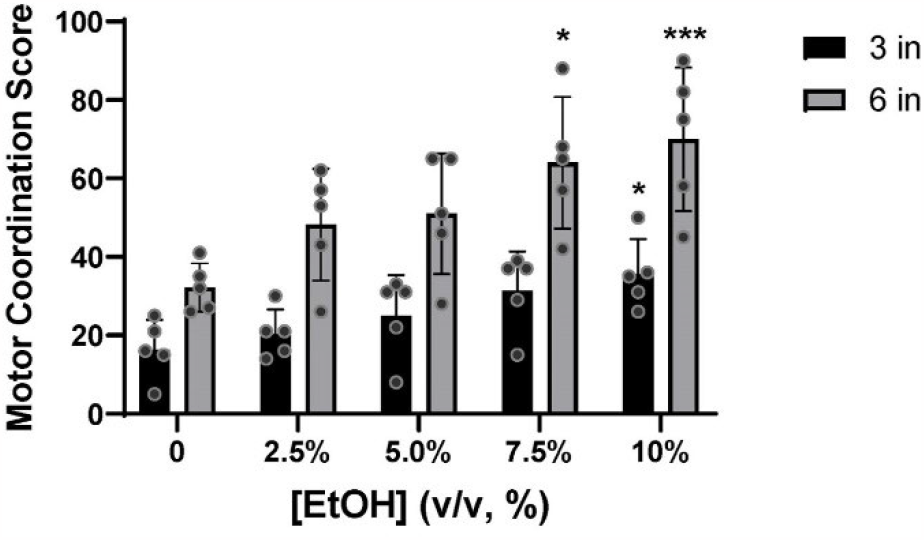
Motor coordination score at 3- and 6-inch height at different ethanol concentrations. At 3 inches, the motor coordination score increased as alcohol concentration increased, with a statistically significant difference relative to baseline being observed at concentrations of 10% (p=0.0201). At 6 inches, the test score showed a similar pattern of increasing and showed significance at 7.5% EtOH (p=0.0277) and 10% EtOH (p=0.0004) compared to baseline (2.5%). Bars represent medians ± 95% CI.

## DISCUSSION

The primary objective of this study was to develop and validate a bespoke HLT apparatus and protocol as a means of evaluating motor coordination deficits in pigs. The data, albeit with a small sample size, are consistent with our predictions. During training, the number of errors observed in the HLT decreased as the rung height decreased. This difference is not attributable to the test being easier to perform at 3 inches, but rather the animals understanding the requirements of the task. This conclusion is supported by the observation of a similar pattern of performance, under varying alcohol concentration exposures, at both 3- and 6-inches rung heights (Figure 5). Giving the short legs and heavy body weight of the pigs, training the animals at a rung height of 6 inches, followed by a gradual decrease to 3 inches across training sessions, is sufficient to train the animals to lift their legs completely when crossing the ladders, minimizing the numbers of errors observed and providing a reasonable baseline measurement of motor coordination.

A secondary objective of this study was to perform an initial evaluation of construct validity of the HLT protocol. Consistent with our predictions, the same animals that were successfully trained on this protocol during the primary objective evaluations, had higher motor coordination scores (i.e., made more errors) as ethanol concentration and consumption to intoxication increased. Together, these results demonstrate that our HST protocol shows good face and construct validity, as it is both suitable for evaluation of motor coordination in pigs as well as sensitive to deficits in motor coordination as a consequence of ethanol consumption.

As an initial evaluation of this HLT protocol, a few issues were observed which warrant discussion. At the group level, the pattern of behavior in motor coordination in the HLT results remain consistent at both 3- and 6-inches as ethanol consumption increased. Nevertheless, individual differences in motor coordination behaviors were observed, likely attributable to individuals’ differences in drinking, both in volume and speed of consumption. This protocol was developed as part of a larger project, which evaluates progression to Alcohol Use Disorder (AUD) at an individual level, and explains our decision to use a free choice alcohol consumption paradigm. It is conceivable, however, that reduced variability would be observed, if a fixed volume and concentration of alcohol were administered via gavage.

The decision to exclusively use female pigs in this study was also influenced by the overarching research question and limitations in terms of available space at our vivarium. Given that this was a pilot study focused on method development and prioritizing investigators’ safety, female pigs were chosen for their generally more docile behavior upon reaching puberty. Correspondingly, male pigs can display agonistic behavior at puberty, particularly when housed in close vicinity to female conspecifics^11,12^. While our decision to study female pigs was therefore pragmatic, we do not predict that sex-differences in motor coordination would be observed in subsequent studies.

The materials for the low-cost, HLT apparatus, sourced from materials common to any big box hardware, affords flexible adjustments of dimensions according to the pigs’ size. By conducting training and testing on flat ground, the risk of injury to the pigs, even when they were heavily intoxicated was minimized. We, therefore, confidently recommend the HLT protocol for use across a wide array of disease models in pigs. This includes applications in the pre- and post-treatment evaluation for substance use disorders including AUD, neurodegenerative diseases such as amyotrophic lateral sclerosis (ALS) and Parkinson’s disease (PD), motor impairments resulting from conditions including multiple sclerosis (MS), strokes, traumatic head injuries, and ataxic disorders of balance, vestibular function and proprioception.

## ACKNOWLEDGMENTS

This study was supported by the Laura W. Bush Institute of Women’s Health, Kayla Weitlauf Endowment for Women’s Health, and NIH AA027401.

## SUPPLEMENTARY VIDEOS

https://figshare.com/s/97c7035297647572d73e

## REFERENCES

1. Paul, S. M. et al. How to improve R&D productivity: the pharmaceutical industry’s grand challenge. Nat Rev Drug Discov. 9 (3), 203–214, doi:10.1038/nrd3078, (2010).

2. Bailoo, J. D., Reichlin, T. S. & Wurbel, H. Refinement of experimental design and conduct in laboratory animal research. ILAR J. 55 (3), 383–391, doi:10.1093/ilar/ilu037, (2014).

3. Gaire, J. et al. Spiny mouse (Acomys): an emerging research organism for regenerative medicine with applications beyond the skin. NPJ Regen Med. 6 (1), 1, doi:10.1038/s41536-020-00111-1, (2021).

4. Ioannidis, J. P. Evolution and translation of research findings: from bench to where? PLoS Clin Trials. 1 (7), e36, doi:10.1371/journal.pctr.0010036, (2006).

5. Chavalarias, D. & Ioannidis, J. P. Science mapping analysis characterizes 235 biases in biomedical research. J Clin Epidemiol. 63 (11), 1205–1215, doi:10.1016/j.jclinepi.2009.12.011, (2010).

6. Ioannidis, J. P. Why most published research findings are false. PLoS Med. 2 (8), e124, doi:10.1371/journal.pmed.0020124, (2005).

7. Goodman, S. & Greenland, S. Why most published research findings are false: problems in the analysis. PLoS Med. 4 (4), e168, doi:10.1371/journal.pmed.0040168, (2007).

8. Regulations, A. W. Animal Welfare Act. Animal Welfare Act. (2013).

9. Lee, J. H. et al. A novel porcine model of traumatic thoracic spinal cord injury. J Neurotrauma. 30 (3), 142–159, doi:10.1089/neu.2012.2386, (2013).

10. Schomberg, D. T. et al. Translational Relevance of Swine Models of Spinal Cord Injury. J Neurotrauma. 34 (3), 541–551, doi:10.1089/neu.2016.4567, (2017).

11. Vodicka, P. et al. The miniature pig as an animal model in biomedical research. Ann N Y Acad Sci. 1049 161–171, doi:10.1196/annals.1334.015, (2005).

12. Lunney, J. K. et al. Importance of the pig as a human biomedical model. Sci Transl Med. 13 (621), eabd5758, doi:10.1126/scitranslmedabd5758, (2021).

13. Murphy, E., Nordquist, R. E. & van der Staay, F. J. A review of behavioural methods to study emotion and mood in pigs, Sus scrofa. Applied Animal Behaviour Science. 159 9–28, doi:10.1016/j.applanim.2014.08.002, (2014).

14. Gieling, E. T., Schuurman, T., Nordquist, R. E. & van der Staay, F. J. in Molecular and Functional Models in Neuropsychiatry (ed Jim J. Hagan) 359–383 (Springer Berlin Heidelberg, 2011).

15. Sullivan, S. et al. Improved behavior, motor, and cognition assessments in neonatal piglets. J Neurotrauma. 30 (20), 1770–1779, doi:10.1089/neu.2013.2913, (2013).

16. Boakye, M. et al. Treadmill-Based Gait Kinematics in the Yucatan Mini Pig. J Neurotrauma. 37 (21), 2277–2291, doi:10.1089/neu.2020.7050, (2020).

17. Shin, S. K. et al. An Adolescent Porcine Model of Voluntary Alcohol Consumption Exhibits Binge Drinking and Motor Deficits in a Two Bottle Choice Test. Alcohol Alcohol. 56 (3), 266–274, doi:10.1093/alcalc/agaa105, (2021).

18. Herbers, J. M. Time resources and laziness in animals. Oecologia. 49 (2), 252–262, doi:10.1007/BF00349198, (1981).

19. Varholick, J. A. et al. Social dominance hierarchy type and rank contribute to phenotypic variation within cages of laboratory mice. Sci Rep. 9 (1), 13650, doi:10.1038/s41598-019-49612-0, (2019).

20. Bailoo, J. D., Jordan, R. L., Garza, X. J. & Tyler, A. N. Brief and long periods of maternal separation affect maternal behavior and offspring behavioral development in C57BL/6 mice. Dev Psychobiol. 56 (4), 674–685, doi:10.1002/dev.21135, (2014).

21. Varholick, J. A., Bailoo, J. D., Palme, R. & Wurbel, H. Phenotypic variability between Social Dominance Ranks in laboratory mice. Sci Rep. 8 (1), 6593, doi:10.1038/s41598-018-24624-4, (2018).

22. Bailoo, J. D. et al. Evaluation of the effects of space allowance on measures of animal welfare in laboratory mice. Sci Rep. 8 (1), 713, doi:10.1038/s41598-017-18493-6, (2018).

23. Bailoo, J. D. et al. Effects of Cage Enrichment on Behavior, Welfare and Outcome Variability in Female Mice. Front Behav Neurosci. 12 232, doi:10.3389/fnbeh.2018.00232, (2018).

24. Bailoo, J. D. et al. Effects of weaning age and housing conditions on phenotypic differences in mice. Sci Rep. 10 (1), 11684, doi:10.1038/s41598-020-68549-3, (2020).

